# DDN2.0: R and Python packages for differential dependency network analysis of biological systems

**DOI:** 10.1101/2021.04.10.439301

**Authors:** Bai Zhang, Yi Fu, Yingzhou Lu, Zhen Zhang, Robert Clarke, Jennifer E. Van Eyk, David M. Herrington, Yue Wang

## Abstract

Data-driven differential dependency network analysis identifies in a complex and often unknown overall molecular circuitry a network of differentially connected molecular entities (pairwise selective coupling or uncoupling depending on the specific phenotypes or experimental conditions) (Herrington, et al. 2018; Zhang, et al., 2009; Zhang and Wang, 2010; Zhang, et al., 2016). Such differential dependency networks are typically used to assist in the inference of potential key pathways. Based on our previously developed Differential Dependency Network (DDN) method, we report here the fully implemented R and Python software tool packages for public use. The DDN2.0 algorithm uses a fused Lasso model and block-wise coordinate descent to estimate both the common and differential edges of dependency networks. The identified DDN can help to provide plausible interpretation of data, gain new insight of disease biology, and generate novel hypotheses for further validation and investigations.

To address the imbalanced sample group problem, we propose a sample-size normalized formulation to correct systematic bias. To address high computational complexity, we propose four strategies to accelerate DDN2.0 learning. The experimental results show that new DDN2.0+ learning speed with combined four accelerating strategies is hundreds of times faster than that of DDN2.0 algorithm on medium-sized data (Fu, 2019). To detect intra-omics and inter-omics network rewiring, we propose multiDDN using a multi-layer signaling model to integrate multi-omics data. The simulation study shows that the multiDDN method can achieve higher accuracy of detecting network rewiring (Fu, 2019).

## 1 Introduction

Data-driven differential network analysis identifies in a complex and often unknown overall molecular regulatory circuitry a network of differentially connected molecular entities (**Fig. 1a**; pairwise selective coupling or uncoupling depending on the specific phenotypes or experimental conditions, including the associated regulatory elements) (Ha, et al., 2015; Herrington, et al. 2018; Hu, et al., 2016; Mitra, et al., 2013; Ottenbros, et al., 2021; Tolios, et al., 2020; Zhang, et al., 2016). Such differential regulatory networks are typically used to assist in the inference of potential key pathways and targets in tumor biology. They can serve as useful frameworks for the construction and verification of mechanistic cancer models, which in turn help to provide plausible interpretation of data, gain new insight of cancer biology, and generate hypotheses for further validation and investigations (Califano, 2011; Gill, et al., 2010; Hudson, et al., 2009; Ideker and Krogan, 2012; Zhang, et al., 2016). It should be noted that the pairwise selective coupling or uncoupling in such data-derived network are based on statistical significance in differential co-expressions, representing possible underlying mechanisms involving direct or indirect, and single or collective effect of a multitude of regulatory pathways.

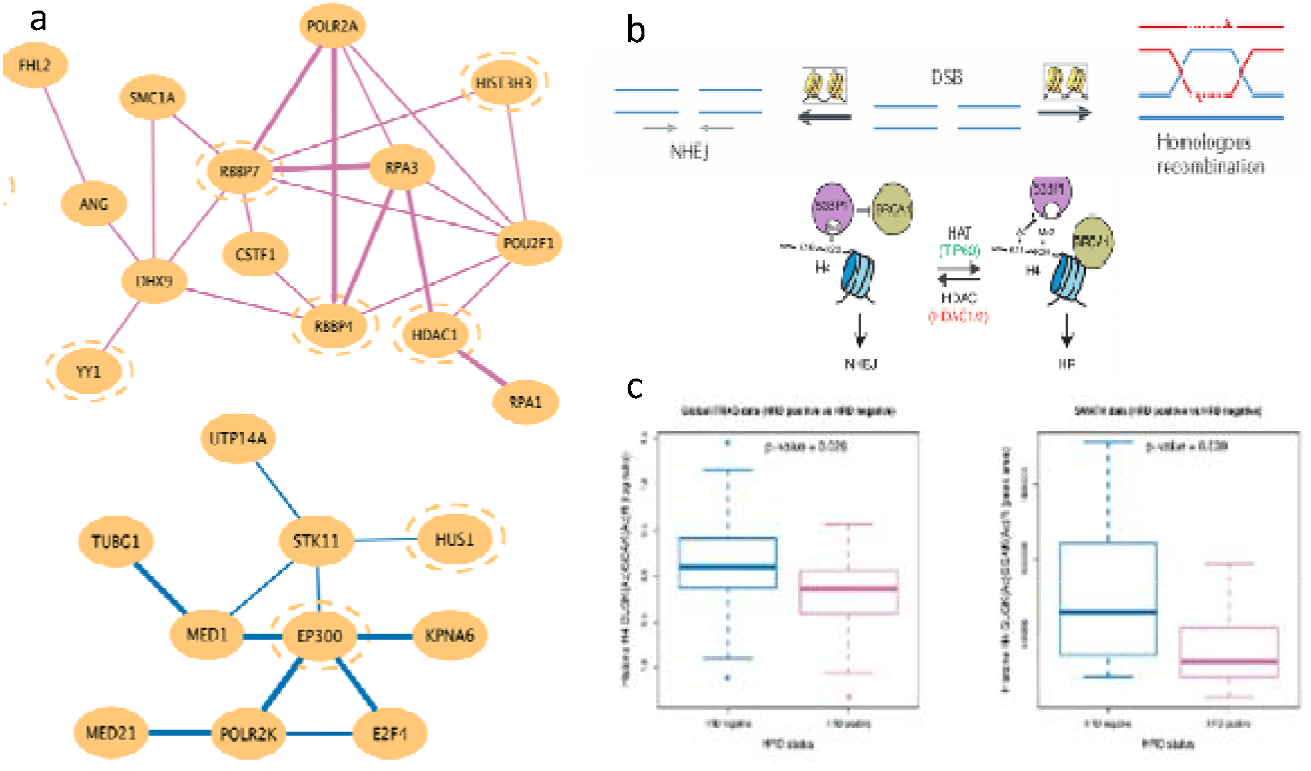

The publicly available Differential Dependency Network (DDN) analysis software tool (Zhang, et al., 2009; Zhang, et al., 2011; Zhang and Wang, 2010) and its recent extension, knowledge-fused DDN (kDDN) (Tian, et al., 2014), represent our continuous effort to develop novel bioinformatics tools to detect statistically significant rewiring and robust subnetworks with phenotype-specific changes. We have successfully applied DDN tools to real gene expression and proteomics data with promising findings and novel hypotheses (Crawford, et al., 2010; Hu, et al., 2015; Tian, et al., 2014; Zhang, et al., 2009; Zhang, et al., 2016). The current DDN tool can infer statistically significant coupling or uncoupling between pairs of molecular entities (e.g., genes/ proteins) that are dependent on the specific phenotypes or experimental conditions using transcriptome/proteome expression data (Tian, et al., 2014; Zhang, et al., 2009; Zhang, et al., 2011). To assess the DDN’s effectiveness, we have compared the accuracy of inferring differential networks by DDN with other tools using realistic simulations (Tian, et al., 2014; Zhang, et al., 2009). The DDN tool has been used to detect critical changes in molecular networks in breast cancer (Madhavan, et al., 2011), ovarian cancer (Zhang, et al., 2016), and medulloblastoma. This has led to novel findings and hypotheses that have been validated experimentally (Hu, et al., 2015; Zhang, et al., 2016).

From a scientific perspective, the main role of DDN is at least two twofold. First, an enrichment analysis focused on the genes/proteins that are differentially connected (nodes of differential networks including associated regulatory elements), not simply differentially expressed, can identify the more relevant pathways that functionally control phenotypes (Barabasi, et al., 2011; Califano, 2011; Hu, et al., 2015; Mitra, et al., 2013). Second, the hubs of differential networks (densely and differentially connected nodes) can hint at changes in the regulatory mechanisms, e.g., PTM or mutations in the regulatory elements of master regulators (Hudson, et al., 2009; Zhang, et al., 2016). By contrast, classic differential or co-expression analysis (i.e., identifying a subset of differentially or co-expressed genes/proteins) is insufficient here because the low level of transcription factors (TF) differential expression makes their detection challenging, even though TFs play a central regulatory role in controlling gene expression (Reverter, et al., 2010). DDN addresses an important biological issue because it better accounts for the functional activation/deactivation of TF (e.g., reversible phosphorylation, enhancer binding, chromatin openness) than does differential expression alone. Simple differential expression will overlook these vital changes in regulatory mechanism. For example, Hudson et al. reported that the myostatin gene containing a causal mutation was not detected because myostatin was not differentially expressed at any of ten developmental time points under surveillance (Hudson, et al., 2009).

The goal of studying network remodeling is not to solve the entire and common network at once but to find small, most-relevant, and critical features (e.g, topological changes within/between pathway networks) that can explain how the network functions to control phenotype. For example, most existing approaches infer gene co-expression networks under a single biological condition and are focused on the disease-related pathways enriched on a subset of differentially expressed genes. In marked contrast, DDN infers differential regulatory networks, i.e., differentially connected genes between different cancer subtypes, and is focused on the disease-related pathways enriched on (i) a subset of multi-omics expressions whose regulatory patterns (e.g., paired miRNA-mRNA/protein co-expressions) meet biological expectations and/or (ii) a subset of differentially connected genes/proteins (selectively coupled or uncoupled protein/gene pairs and associated regulatory elements depending on the specific cancer subtype). Another advantage is that we obtain small/sparse experimentally tractable models, i.e., rather than complex hairballs, DDN generally produces models that allow for the design of feasible wet lab experiments that enable mechanistic validation (Hu, et al., 2015; Tyson, et al., 2011; Zhang, et al., 2016).

## 2 Results

### Case study on initial application of DDN to CPTAC proteomics data

In our recent work on integrated proteogenomic characterization of human high grade serous ovarian cancer (Zhang, et al., 2016), applying DDN on a set of 171 BRCA1/2-related proteins, we identified a sub-network of 30 proteins that displayed differential co-expression patterns between HRD (homologous recombination deficiency) from non-HRD patients (**Fig. 1a**). Several of the proteins in these modules are known to be involved in histone acetylation or deacetylation. Although statistical association predicted by DDN cannot distinguish between drivers and consequences of HRD status, the observed enrichment of proteins associated with histone acetylation motivated us to identify and quantify acetylated peptides from our CPTAC global proteomic data. Comparative analysis of 399 acetylated peptides identified 15 acetylated peptides with significant differences between the HRD and non-HRD samples (dual acetylation at K12 and K16 of histone H4). Acetylation of H4 is involved in the choice of DNA double strand break (DSB) repair pathways (homologous recombination or non-homologous end joining). This relationship is regulated partially by HDAC1 (**Fig. 1b**). The findings were verified using independent samples and independent technical approaches (**Fig. 1c**) (Zhang, et al., 2016). The combined observations of increased HDAC1 (a hub protein also identified by DDN) and associated proteins at the pathway level, together with decreased acetylation of H4 in HRD patients at the (post-translational modifications) PTM level, provide insight regarding the potential role of HDAC1 in modulating the choice of DSB repair pathways (Zhang, et al., 2016).

### Case study to validate phenotype-specific differential networks of breast cancer dormancy and/or resistance

We have been previously funded to study how gene networks may adapt to affect the biology of breast cancers. Our CCSB/CSBC project is focused on understanding how estrogen receptor-positive (ER+) breast cancer cells adapt to the stress of endocrine-based therapies. Our central hypothesis invokes a gene network that coordinately regulates the functions of a cell and determines and executes the cell’s fate decision. Using DDN tool we were able to identify three small topological features, one of which independently reflected much of our prior knowledge despite not explicitly incorporating this knowledge in learning the models (Zhang, et al., 2009). We followed the predictions of this <u>differential topology</u> and validated fundamentally new insights into molecular signaling, e.g., direct regulation of BCL2 by XBP1 and the requirement of NFκB for XBP1 signaling to regulate the pro-survival cell fate outcome in the cellular context of antiestrogen treatment and resistance (E2 versus E2+ICI) (Fig. 2) (Clarke, et al., 2011; Crawford, et al., 2010; Hu, et al., 2015; Tyson, et al., 2011). Specifically, DDN tool identified a new non-canonical UPR signaling route that was not previously known. Canonical signaling in the UPR would have predicted that changes in JNK regulated BCL2 but DDN predicted that XBP1 (not JNK) regulated BCL2 (Zhang, et al., 2009). We then found (i) response elements to XBP1 in the BCL2 promoter, (ii) that cells over expressing endogenous XBP1 also had higher BCL2 expression, (iii) that knockdown of XBP1 in these cells with RNAi reduced BCL2 expression, (iv) that constitutive overexpression of the XBP1 cDNA increased BCL2 expression and (v) that knockdown of BCL2 phenocopies knockdown of XBP1 with respect to reversing endocrine resistance (Crawford, et al., 2010; Hu, et al., 2015). We went on to further explore the DDN model and further confirmed the role of XBP1 regulation of NFkB (also not previously known) as another key driver of the prosurvival activities of XBP1 and the UPR (Hu, et al., 2015). In more detail, XBP1 can function as a direct/co-regulator through binding to its responsive element, most notably ERα. BCL2 contains response elements for both ERa and XBP1, thus XBP1-BCL2 may either be rewired or involve ERa as a latent variable, or intervening gene. In the non-recurring breast cancer, the affected network involves both signals received from activation of the membrane receptors and a cascade of signaling path inside a cell to promote apoptosis as well as survival. A balance between apoptosis and survival is necessary for damaged cells to be eliminated and repaired cells to survive. Three immune response genes (IL1B, NFκB and TNFα) for increased resistance to breast cancer treatment were identified in the recurring tumors. These genes formed a path to inhibit pro-apoptotic CASP3 and PPP3R1, and to activate pro-survival gene PIK3R5 or CSF2RB.

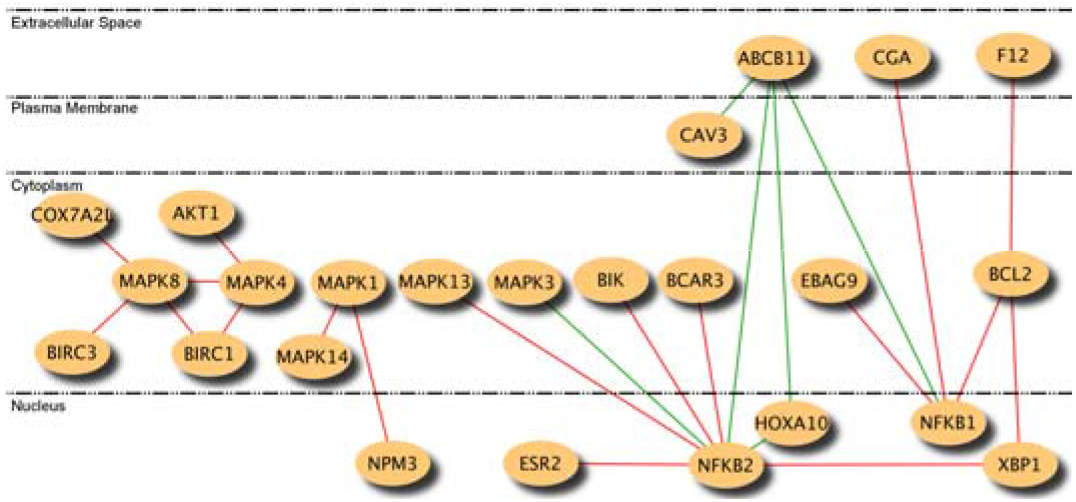

### Case study to validation of DDN prediction on CPTAC data and identify phenotype-associated co-regulatory subnetworks

In our recent work (Zhang, et al., 2016), we reported a key finding on proteogenomic analysis of TCGA ovarian tumors in which DDN played a critical role (**Fig. 1**). The differentially networked proteins identified by DDN analysis on global proteomics data suggest that histone acetylation might play a role in DNA repair. We studied the downstream effects and identified several peptides of histone H4 showing differentially acetylated K12 and K16. We validated these discoveries using SWATH proteomics technology, in addition to a separate yet direct evidence demonstrated on a cell line model. In more detail, DDN analysis on ovarian cancer samples helped identifying a sub-network of 30 proteins involved in histone acetylation or deacetylation (**Fig. 1**). In cell line data, acetylation of H4 has previously been reported to be involved in the choice of DNA double strand break (DSB) repair pathways (homologous recombination or non-homologous end joining). This relationship is regulated partially by HDAC1, a protein also identified in the DDN analysis. We observed a significant enrichment of HDAC1 and its co-regulated proteins in tumors with HRD and low H4 acetylation relative to non-HRD tumors with high H4 acetylation (DDN analysis: permutation tests, p value <0.05; differentially acetylated peptides: t-tests, p value <0.05 with an estimated FDR < 0.5% by bootstrap/permutation tests). The identification of these acetylation events associated with HRD may provide an alternative biomarker of HRD for patient selection in future clinical trials of HDAC inhibitors alone or in combination with PARP inhibition. This may help to resolve the current discrepancy between the initial observation of limited single agent activity of HDAC inhibitors in ovarian cancer and more recent findings of a >40% response rate when used in combination with cytotoxic chemotherapy in platinum-resistant patients (Zhang, et al., 2016).

The integration of proteogenomic data also refers to the integration of molecular networks across different phenotypes/conditions to identify functional modules that are rewired in response to conditional fluxes. The types of networks themselves could be physical interaction networks, signaling and metabolic pathways, and others. The inference of conditional network often requires the ability to bridge between prior knowledge and experimental data and with built-in resampling/permutation to offer statistically sound and robust results (Zhang, et al., 2014). It should be noted that while DDN infers both common and differential networks, its unique and main function is to differentially detect the statistically significant “rewiring” associated with phenotypic differences. Thus, only rewired edges are included in the final differential dependency networks that are not meant to represent the complete regulatory network that supports the entire functions of the phenotypes. By definition, the rewired part of the networks are not supposed to be huge. We emphasize that the entire molecular machinery behind HR repairing of DSB is much more completed than the DDN detected. However, <u>this is the part that contains key elements</u> differentiating HRD and non-HRD and pin-pointed to histone H4 peptide acetylation as a critical point of the mechanism.

### Case study to analyze PTM interactome

The aforementioned findings of HRD status-dependent H4 acetylations, consistent with recent reports in the literature, indicate that much of the cancer associated molecular alterations will not necessarily be limited to the form of coordinated variations among DNA, copy number, and mRNA/protein expression, but rather as complex rewiring of signaling pathways/networks carried out in the protein and post-translational modification (PTM) interactomes. The DDN in its current form, and more so with its proposed addition, can play an important role in differential analysis of PTM interactomes to capture phenotype/experiment condition-dependent changes. As an example, through CPTAC, we analyzed the tyrosine phosphorylation substrate activity data using the HuProt protein arrays(Hu, et al., 2014). The LC-MS/MS global protein profiles include those proteins that are known pTyr kinases. DDN was applied to identify subnetworks of highly correlated kinase (MS data) – substrate (HuProt array data) pairs that are dependent on HRD phenotypes. The identified subnetworks were further compared against the previously experimentally generated kinase-substrate relationship (KSR) map(Hu, et al., 2014) to retain only connections that are known possible KSRs (**Fig. 3a**). We were able to identify several kinase hubs, such as MAP2K4. Many of them are known drug targets (unpublished, on-going research) (**Fig. 3b**). In this example, the relationship between K and S nodes are directional (downstream substrates). The superposition of KSR map was applied separately. The extension of DDN to allow for seamless integration of such knowledge in a quantitative and computational way will further optimize such analysis approaches.

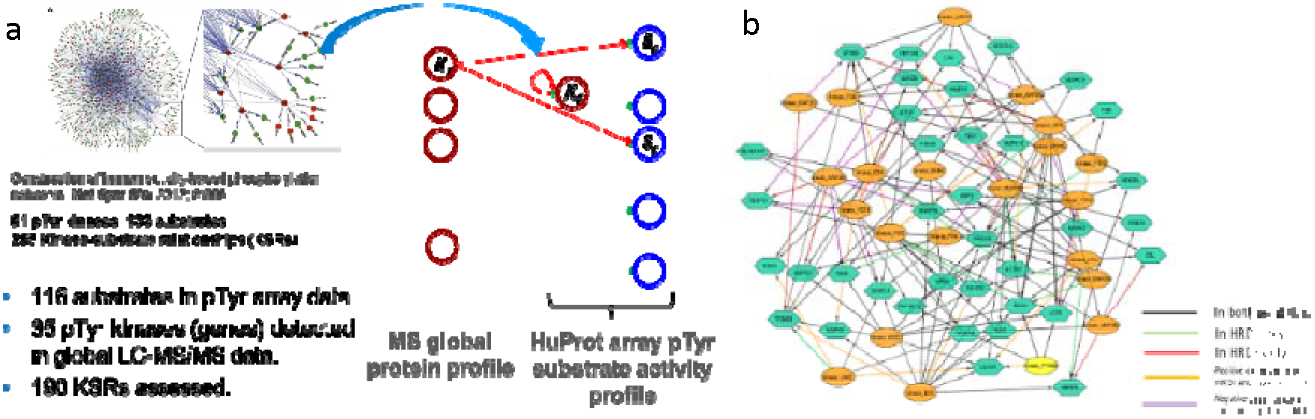

### Case study on cancer prevention research

DDN tool has generated biologically meaningful hypotheses that guided further investigations. To study the consequence of possible transcriptional re-programming regulated by promoter methylation status (effectors ER, BCL2, LEP, and EGR1; AKT1 can regulate methylation patterns in some promoters, *e.g*., AKT1-EGR1 edge), revealed by DDN, we found that *in utero* estrogen exposures induce a rewired network pattern in the mammary glands of rodent offspring that predicts for resistance to endocrine therapies. Subsequent studies have shown that tumors that arise in these mammary glands are less responsive to Tamoxifen (TAM) (Hilakivi-Clarke, et al., 2017). <u>This represents the first study to explain why many ER+ breast cancers fail to respond (or respond and later recur) with TAM treatment</u> (**Fig. 4**).

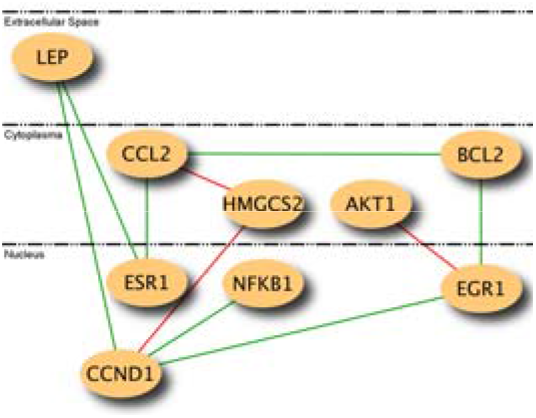

## 3. DDN2.0+ new development

### 3.1 Correction of imbalanced sample size between two groups

Though DDN2.0 is a powerful differential network analysis tool (Zhang, et al., 2009; Zhang and Wang, 2010), some drawbacks hinder the DDN tool from being more widely used in network study or biomedical data analysis. Firstly, the initial design of the DDN method is for comparing two groups with equal sizes of samples. In practice, we notice that for imbalance data, i.e., two groups have different sample sizes, DDN seems to detect network rewiring favoring one condition over the other. A systematic bias caused by data imbalance may exist and need to be correct. Secondly, when limited by the algorithm running time, DDN can handle only dozens of features and samples. The actual computation time grows more than cubically with a feature scale. In biological network analysis, the feature size in gene regulatory network inference could be as large as tens of thousands. The development of accelerated DDN learning algorithms is an urgent need for extending DDN’s application to broader fields. Thirdly, for omics data analysis, the current DDN method is designed for analyzing a single type of omics data. Though a simple method of merging multiple data matrices into a single matrix could theoretically extend the DDN method to multi-omic data, this kind of integration method would double to triple the feature scale and take impractically long computation time. This simple merging method also ignores inter-omics regulation knowledge. We believe a multi-omics integration method could take advantage of the additional knowledge such as directional inter-omics interactions (Buescher, et al., 2016), to effectively reduce the feature scale and increase the accuracy in differential network learning. With the advent of more are more multi-omics research projects, DDN needs such an integration method to discover novel network rewiring facilitated by multi-omics data.

In detecting network rewiring between two conditions, we expect that network rewiring events are sparse. In other words, we assume that the networks under two conditions will share a large portion of common network structures. Based on this assumption, Zhang and Wang (2010) further improved the initial DDN method by jointly solving Fused LASSO regressions via introducing a penalty term on the structural difference. We denote this version of the DDN method as DDN2.0. The DDN2.0 optimization problem is:

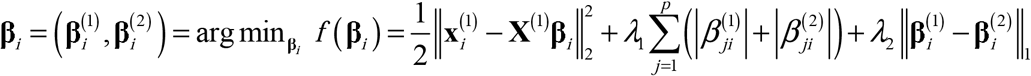

where *i* is the node index; *β*_*ji*_ is the regression coefficient from node *i* to node *j* under a specific condition; ***y***_*i*_ and ***X*** are the expression values of dependent and input variables, respectively; *P* is the number of nodes; *λ*_1_ and *λ*_2_ are the parameters on the two penalty terms that are used to assure both a sparse common network structure and sparse differential network rewiring.

The differences between 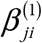 and 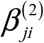 indicate the differential dependence edges, while common dependence edges in the network are inferred by the consistent coefficients. Note that the differential dependences are of particular interest, because such network rewiring may reveal pivotal information on how the biological system responds to different biological conditions. The permutation test is introduced in DDN to evaluate empirical p-values of the detected network rewiring (Tian, et al., 2011). The detected differential edges with multi-test corrected p-values less than the preset significance threshold (e.g., p-value<0.05) are marked as significant network rewiring.

However, in the design of the DDN framework, the objective function does not explicitly consider the sample size difference between the two groups. If we expand the objective function of DDN, we have:

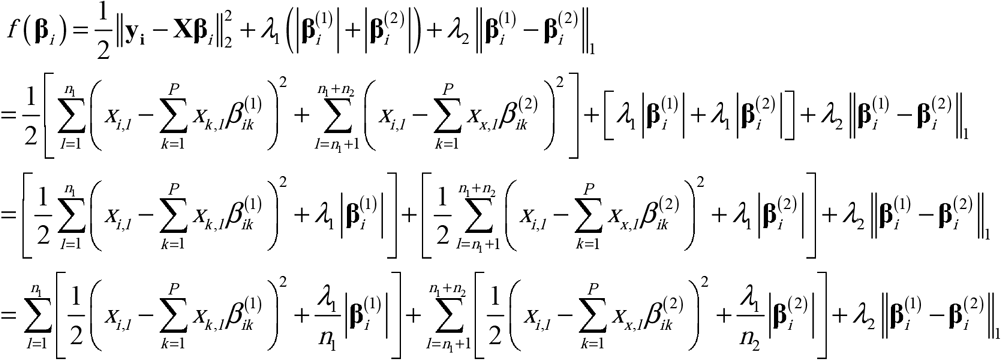

We can see that, although the weights for each group (case vs. control) are equal, and the sample-wise squared residual is at a comparable level, the actual weighted penalty of beta added to each sample differs. Suppose one group’s sample scale is ten times as the other group, the actual sample-wise weight penalty term is as small as one-tenth of the other group. When minimizing the total object function with the L-1 penalty, the samples in the group of smaller sizes would be more likely to have smaller sparsity in network structure.

To redesign the weights applied to each group, we first define two LASSO objective functions(Friedman, et al., 2017) for each group, and rewrite the DDN objective function as follows:

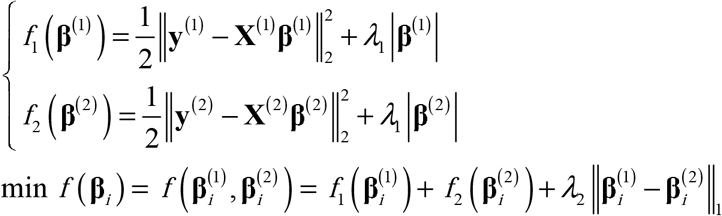

in which *f*_1_ only contains measurement and regression coefficients of condition 1, and so does *f*_2_ to condition 2.

For standardized data, the LASSO objective functions *f*_1_ and *f*_2_ is positively dependent on the sample size N. To make the two group’s objective function values at a comparable level, we simply add sample scale normalizer to the basic form of LASSO objective function. Define the LASSO objective function with the sample scale normalizer as:

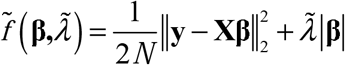

where 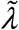 is the parameter for controlling the sparsity. Basically, we use mean squared error to replace the square error term in the original LASSO objective function. This normalized LASSO objective function is the scaled version of basic LASSO form, with 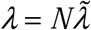:

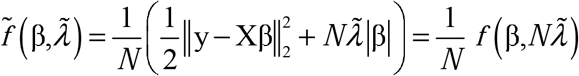

Therefore, the solutions to these two forms of LASSO objective functions are identical, with the condition of 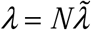. When the value 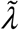 is given, the normalized LASSO objective function is independent of the sample scale N. We now adjust DDN’s formulation accordingly to the normalized LASSO objective function, and hence make the new DDN objective function also independent of the sample scales. The new DDN objective function is:

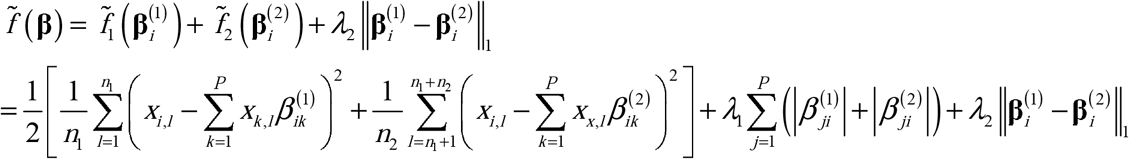

We designed a simulation experiment to confirm the bias brought by imbalanced data in the original DDN method, and also to show our proposed reformulated DDN objective function is capable of handling imbalanced data correctly. To illustrate the systematic bias, we need to identify which differential edges are the false positive detections caused by the bias. We purposely designed the data that contains no groud truth positives of differential edges, and hence all detected differential edges will be false positives. The simulated data is actually single condition data that follow i.i.d multi-variate Gaussian distribution, and is manually divided into two groups to form the pseudo two conditions. We generate data with the total sample scale N=500 and the feature scale P=30, and then divided into two groups with three settings of sample ratios: 1:1, 1:10, and 10:1. The first sample ratio of 1:1 represents the perfect balance of sample scale, which corresponds to balanced data; the rest two sample ratios correspond to imbalanced data wither the larger sample scale in either the first or the second group.

The generated data are standardized to zero-mean and unit variance, and then test by the original DDN method and the reformulated DDN method. The original DDN method is performed by kDDN plugin in Cytoscape without knowledge input, which is mathematically equivalent to the DDN2 method. The reformulated DDN method is implemented by the R language. Its results are visualized by Cytoscape to compare with results from the original DDN method.

The simulation results are shown Figure 5. The condition-specific differential edges are colored as red or green for condition 1 or 2. The edge width is mapped from p-values evaluated by a permutation test, and wider edges have smaller p-values. For data group with a sample size ratio of 1:1, which is balanced data, the original DDN method and the sample-scale-normalized DDN method gave identical results: only two network rewirings events with insignificant p-values are detected. These results fit the expectation since there should be no network rewiring in the simulated data. For imbalanced data groups with sample size ratios of 1:10 and 10:1, original DDN2.0 gives network rewiring detection results of overwhelmingly one-sided condition-specific network rewiring. Some of the detected differential edges have significant p-values evaluated from the permutation test. Since there is no network rewiring in the designed ground truth network, all these condition-specific network rewiring detected by the original DDN2.0 method are false positives, and they confirm the existence of systematic bias brought by imbalanced data. On the other hand, the sample-scale-normalized DDN2.0+ method detected only one or zero network rewiring events in these imbalanced data. The detected differential edges have insignificant p-values and could be further filtered. The results show the low false positive rate of the sample-scale-normalized DDN2.0+ method in the simulation data. The systematic bias in the original DDN2.0 method, i.e., the condition-specific false positives in the imbalanced data, has been successfully corrected.

**Figure 5.**
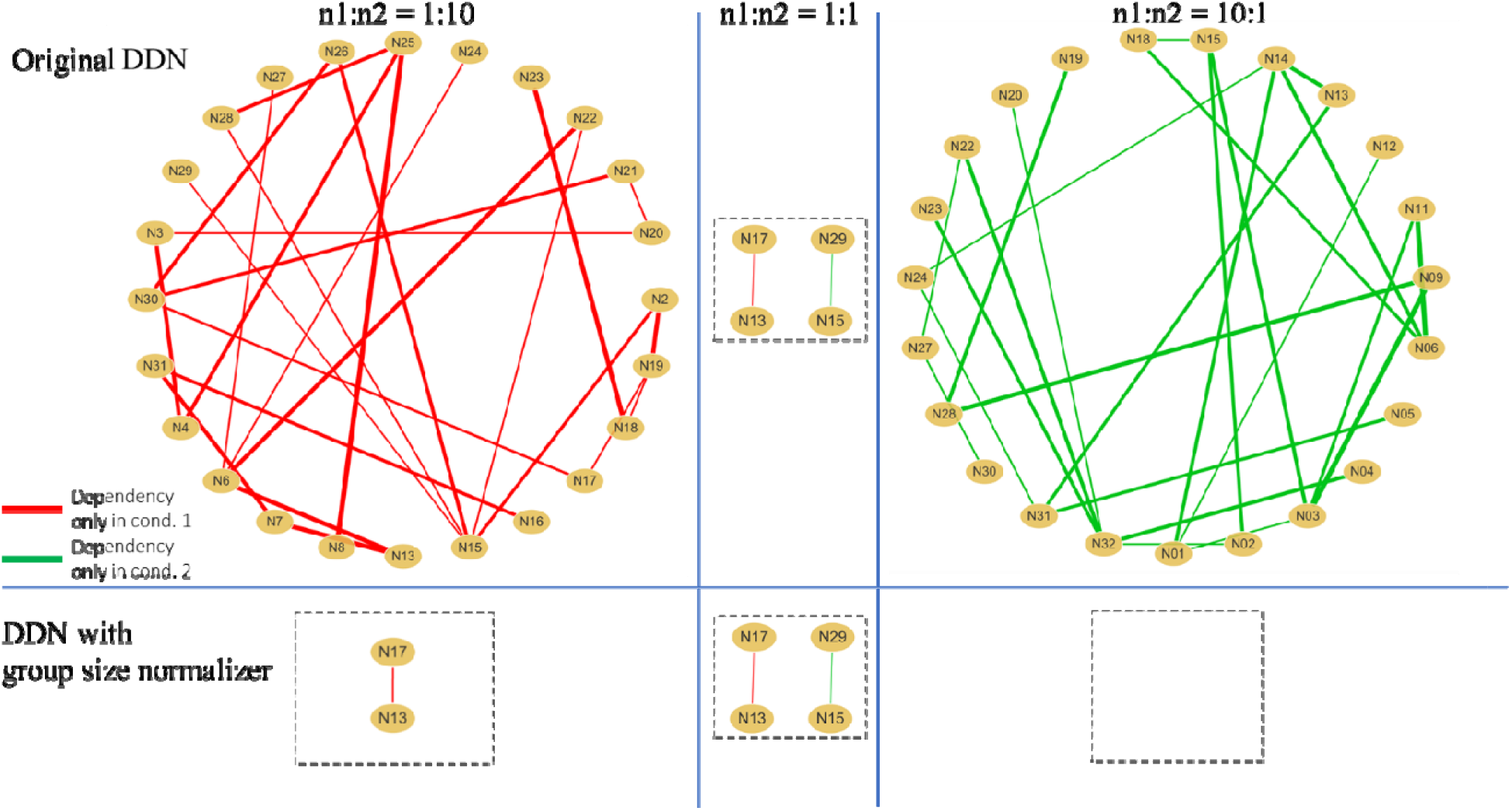
Comparison of DDN2.0 vs DDN2.0+ detected network rewiring from data with different group size ratios.

### 3.2 Accelerated DDN2.0+ algorithm

#### 3.2.1 BCD ResiUpd strategy

We rewrite the DDN2.0+’s objective function into two parts as follows:

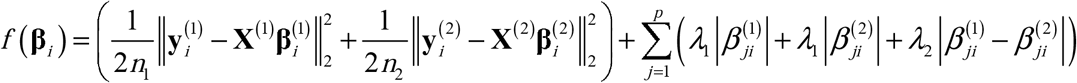

The second part of the objective function can be written as the sum of *p* terms with non-overlapping members 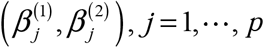. Each 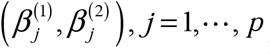 is a coordinate block. The essence of the BCD algorithm is “one-block-at-a-time”. At iteration r + 1, only one coordinate block, 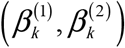 is updated, with the remaining 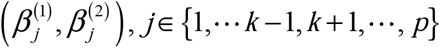 fixed at their values. The cyclic rule is used to update parameter estimation iteratively, i.e., update parameter pair 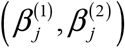 for each *j* = 1, ⋯, *p*, one by one in a circular way in the iterations.

In DDN2.0+, we will derive the solution of 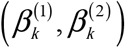 that updated at the end of each iteration. Take DDN objective function’s partial derivative to 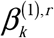 and 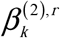:

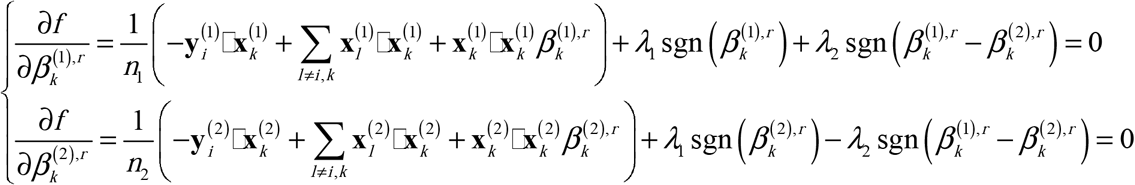

Define the residual as:

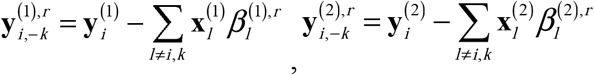

And define the inner products between the residuals and current node’s observation:

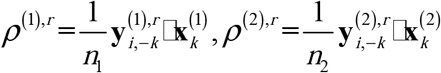

Recalling that the standardized data have unit-variance, the partial derivative equations could be further written as:

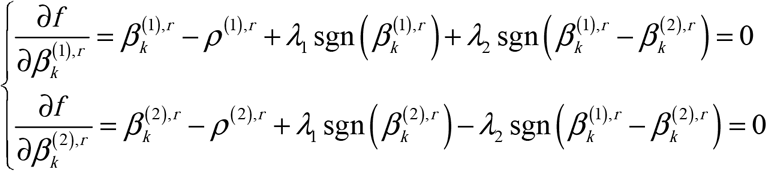

Giving the sign of 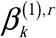, 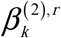 and 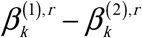, we can get closed-form solutions of 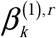 and 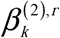. For example, when the conditions are 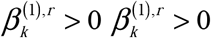 and 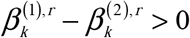, the solution is

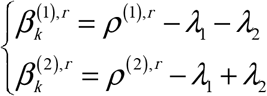

And the condition could then be converted to a sub-region in the plane of (*ρ*^(1)*r*^, *ρ*^(2)*r*^), which is:

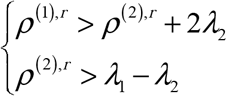

Similarly, we can get all other closed-form solutions for possible conditions, and convert the conditions accordingly to sub-regions on the plane of (*ρ*^(1)*r*^, *ρ*^(2)*r*^). We list all solutions and corresponding subregions as follows

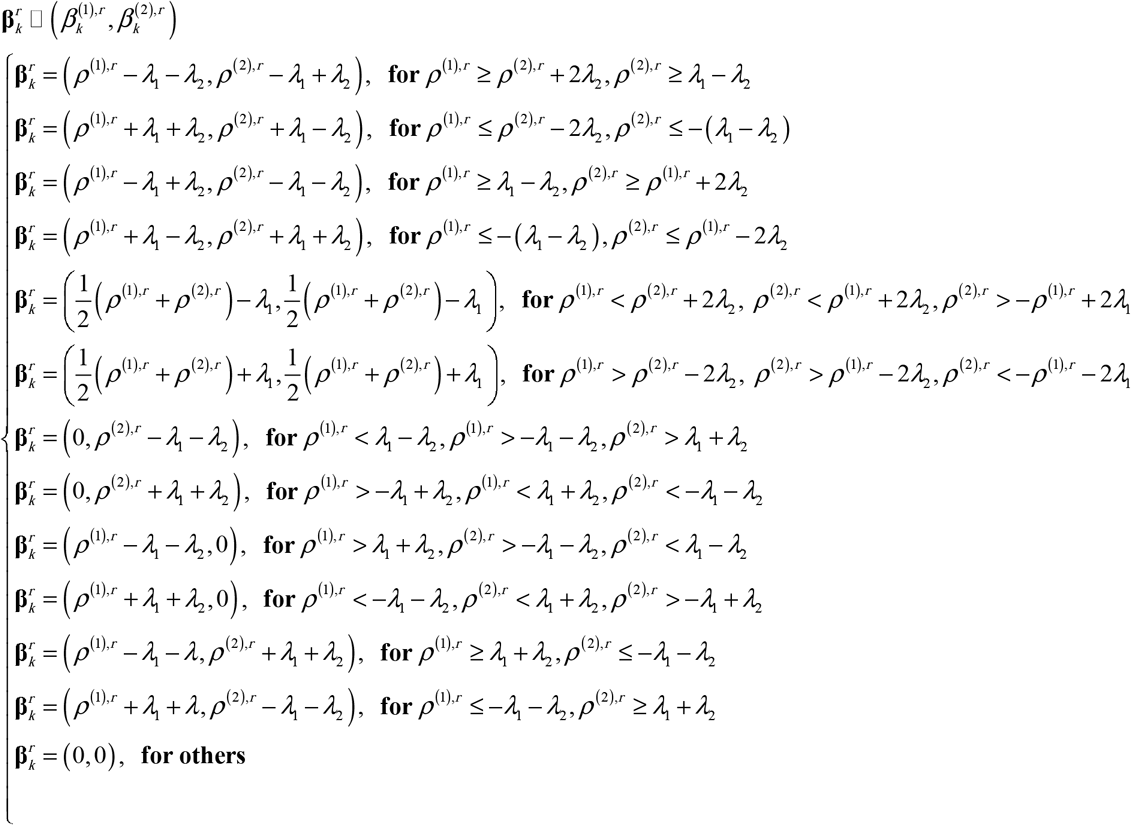

In Figure 6, we illustrate the solution subregions of 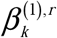 and 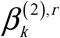 on the plane of (*ρ*^(1)*r*^, *ρ*^(2)*r*^). Recalling that in the BCD algorithm we only update one coordinate at a time, the updated **β**^*r*^ will has most of its elements overlapped with those of **β**^*r*−1^ from the previous iteration. Reinspection on the residuals 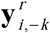 used in the original form of *ρ*^(1)*r*^ or *ρ*^(2)*r*^ definition reveals its relationship with the previous residual:

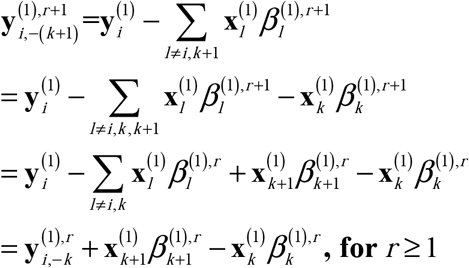

**Figure 6.**
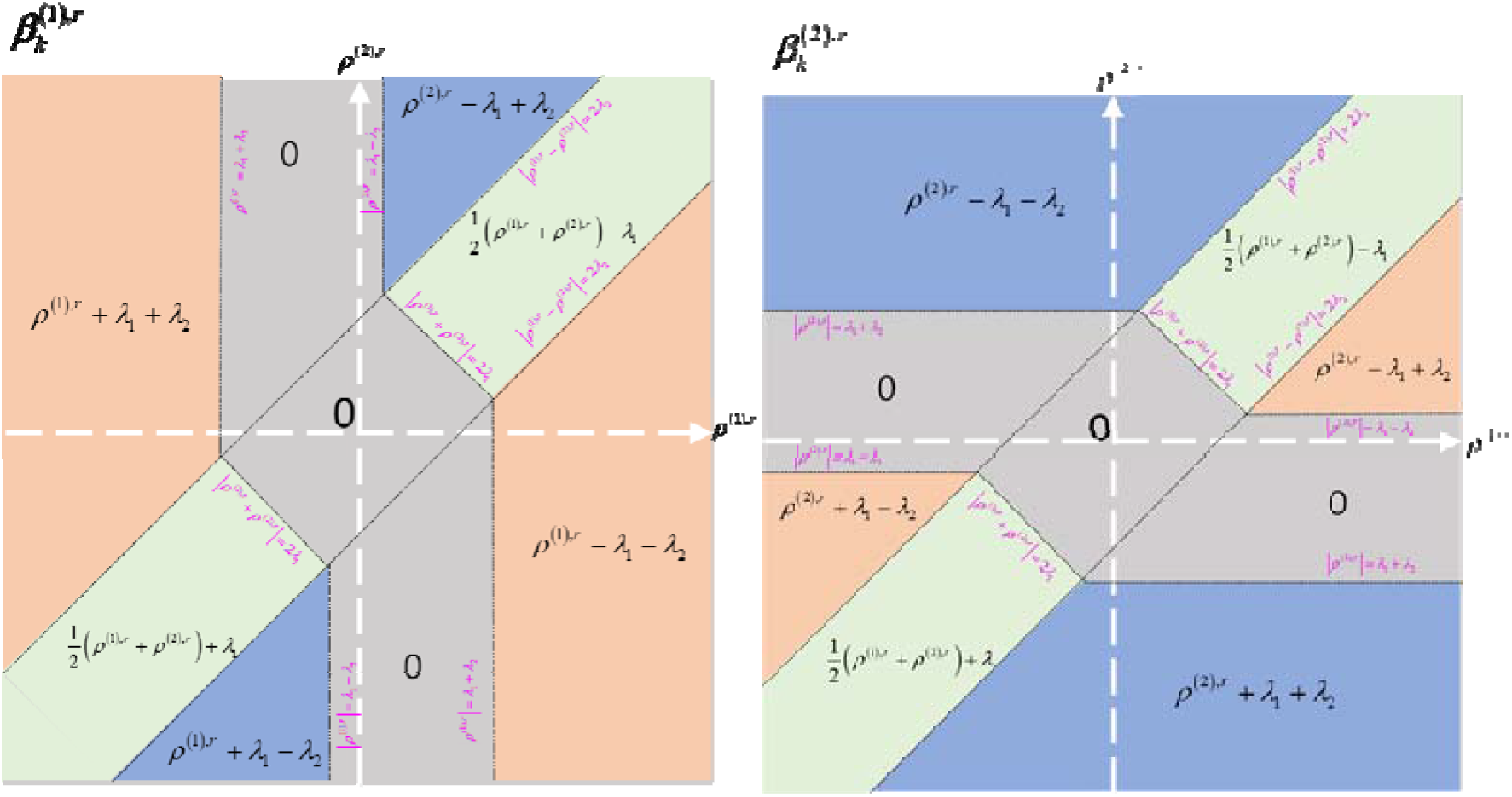
BCD solution subregions on the plane of rho1 and rho2.

Instead of directly calculating residuals from P weighted observed vectors, we can update the residuals in a new iteration from the previous one, with a small computation load on adding 2 weighted observation vectors. We call this computation skill as the method of BCD with residual updating, or BCD-ResiUpd algorithm, and notated the DDN method utilizing this BCD-ResiUpd algorithm as DDN-ResiUpd.

#### 3.2.2 BCD-CorrMtx strategy

In the original form of *ρ*^(1)*r*^ and *ρ*^(2)*r*^, it firstly needs to calculate the residuals 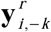 which are the summation of P weighted observation vectors, and takes about PN times of multiplication to complete. Second, the inner product of the residual vector and the current response variable vector requires another N time of multiplication operation. Therefore, calculation of *ρ*^(1)*r*^ and *ρ*^(2)*r*^ in each BCD update needs about 2 ×(*PN* + *N*) ≅ 2*PN* times of multiplication. And in conclusion, the computation complexity of DDN is about *O*(*TP*^2^ × 2*PN*) = *O*(2*P*^3^*N*). For observations with large sample scale N or feature scale P, it will be a heavy burden of computation.

Consider the fact of zero-mean and unit-variance for the standardized data, the inner products between two observed data vectors are actually linear to the elements in their covariance matrix, or equivalently, the Pearson’s correlation matrix R. Therefore, we can replace the inner product of observed data by the pre-calculated correlation coefficients: 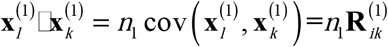 and 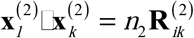. In the neighborhood selection approach, the response variable is one of the nodes: 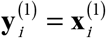. Recalling the definition of *ρ*^(1)*r*^ and *ρ*^(2)*r*^, We rewrite them in the form of correlation matrix elements:

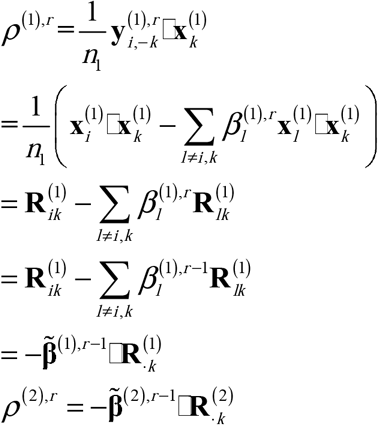

In which

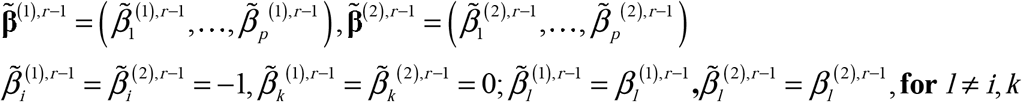

Therefore, *ρ*^(1)*r*^ and *ρ*^(2)*r*^ could be directly calculated from **β**^*r*−1^ and the observation’s correlation matrix R^(1)^ and R^(2)^, without the need for direct computing on the original data matrix X. We call this computation skill as the BCD algorithm using the correlation matrix, or BCD-CorrMtx for short, and notated the DDN method utilizing this BCD-CorrMtx algorithm as DDN-CorrMtx.

The new form of *ρ*^(1)*r*^ or *ρ*^(2)*r*^ has only one inner product of two P-element vectors, and thus needs only 2P times of multiplication operations in each updating iteration. Therefore, the computation complexity of DDN-Corr is *O*(*TP*^2^ × 2*P*) = *O*(2*TP*^3^), approximately N times faster than the procedure with the original definition.

#### 3.2.3 BCD StrongRule strategy

Tibshirani, et al. (2012) proposed “Strong” rules for discarding predictors in Lasso-type problems before for computational efficiency. The Strong rules are developed based on the “Safe” rules proposed by El Ghaoui et al. (2010), and are able to effectively reduce the actual number of predictors need to be solved in LASSO problems. Tibshirani, et al. (2012) also showed that, although in extremely rare cases the Strong rules may erroneously discarding predictors, the error could be amended by checking the KKT conditions. The basic Strong rule is defined as follows: for the lasso problem, discard the j-th predictor from the optimization problem if:

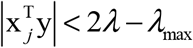

where 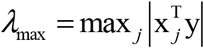 is the smallest tuning parameter value such that 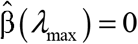.

For the BCD algorithm used in DDN2.0, we use a cross-validation strategy to estimate the optimal value of *λ*_1_ while setting *λ*_2_ = 0. Because DDN2.0 optimization is degraded to basic LASSO problems in this case, DDN2.0+ applies the “Strong” rule to discard predictors before applying the BCD algorithm. The remaining predictors in DDN2.0+ after applying the basic Strong rule will still cover all true predictors, and the results will be identical to predictors without applying the Strong rule. Since the number of remaining predictors is much smaller than the total number of matrix elements, the procedure of LASSO problem solving in DDN2.0+ will take much less computation time (Tibshirani, et al., 2012).

#### 3.2.4 BCD with parallel computing

In the procedure of the neighborhood selection approach, the solution of one node’s optimization is independent of the other nodes’ solution. Therefore, the BCD optimization in DDN2.0+ could work parallelly for each node selected from the total P nodes in the network. In parallel computing, we assign one CPU core from a multi-core computer to independently solve one node’s DDN optimization problem. The whole DDN network construction could be about *N_core* times fast, in which *N_core* is the number of available CPU cores. We developed an R package to implement DDN2.0+ parallel computing along with other accelerating methods of DDN2.0+-CorrMtx or DDN2.0+-ResiUpd.

#### 3.2.5 Simulation studies

We set a series of simulation studies to compare the actual computation time of DDN2.0+ with the four proposed accelerating strategies: DDN-CorrMtx, DDN-ResiUpd, DDN-StrongRule and DDN-CorrMtx/ResiUpd-Parallel. To compare the methods on a fairground, we set the simulation conditions as follows: each of the simulated P nodes’ observation data follows i.i.d standard Gaussian distribution; the true covariance matrix is set to the identity matrix, hence no edges exist in the graph and only the first round of iterations is needed for each node; *λ*_1_ and *λ*_2_ are also set to big enough, therefore the convergence is expected to achieve after the first round of iterations. For each case of simulated data, all methods are using the same data as inputs. The testing environment is listed as follows: CPU Intel® Core™ i5-8300H @2.30Ghz; RAM 16GB; R version 3.6.1. The computation time is recorded by the R function of *system.time()*.

Table 1 listed two simulated cases: one with high feature scale P and the other with high sample scale N. From the comparison of computation time used for each method we could see that, the original DDN method takes the longest time which is unbearable in large scale; DDN with correlation matrix takes little time for large sample scale N, but is slower than DDN with residual updating strategy in case of large feature scale P. Parallel computing offers great help when P is large, but the time saving is limited in case of moderate value of P.

**Table 1.** The computation time of accelerated DDN2.0+ methods

|  | N=100, P=1600 | N=1600, P=100 |
| --- | --- | --- |
| Original DDN | 2695.48s | 141.25s |
| DDN-CorrMtx | 93.19s | 0.28s |
| DDN-CorrMtx-Parallel (nCore=7) | 1.48s | 0.25s |
| DDN-ResiUpd | 31.39s | 2.94s |

**Table 2.** Computation time comparison between DDN2.0+ with and without Strong rule

|  | N=100, P=400 | N=100, P=800 |
| --- | --- | --- |
| DDN-CorrMtx | 57.86s | 393.15s |
| DDN-CorrMtx+<br>Strong Rule | 2.15s | 5.91s |

Figure 7 shows the computation time used by DDN-CorrMtx, DDN-ResiUpd and the original DDN versus the feature scale P. The DDN-ResiUpd method shows a large advantage when P grows. ***Figure*** shows the computation time used by the three methods versus sample scale N. Since the size of the correlation matrix only relies on P, the DDN-CorrMtx method uses almost the same time for different sample scales, and hence is much faster than the other two methods when N is large.

**Figure 7.**
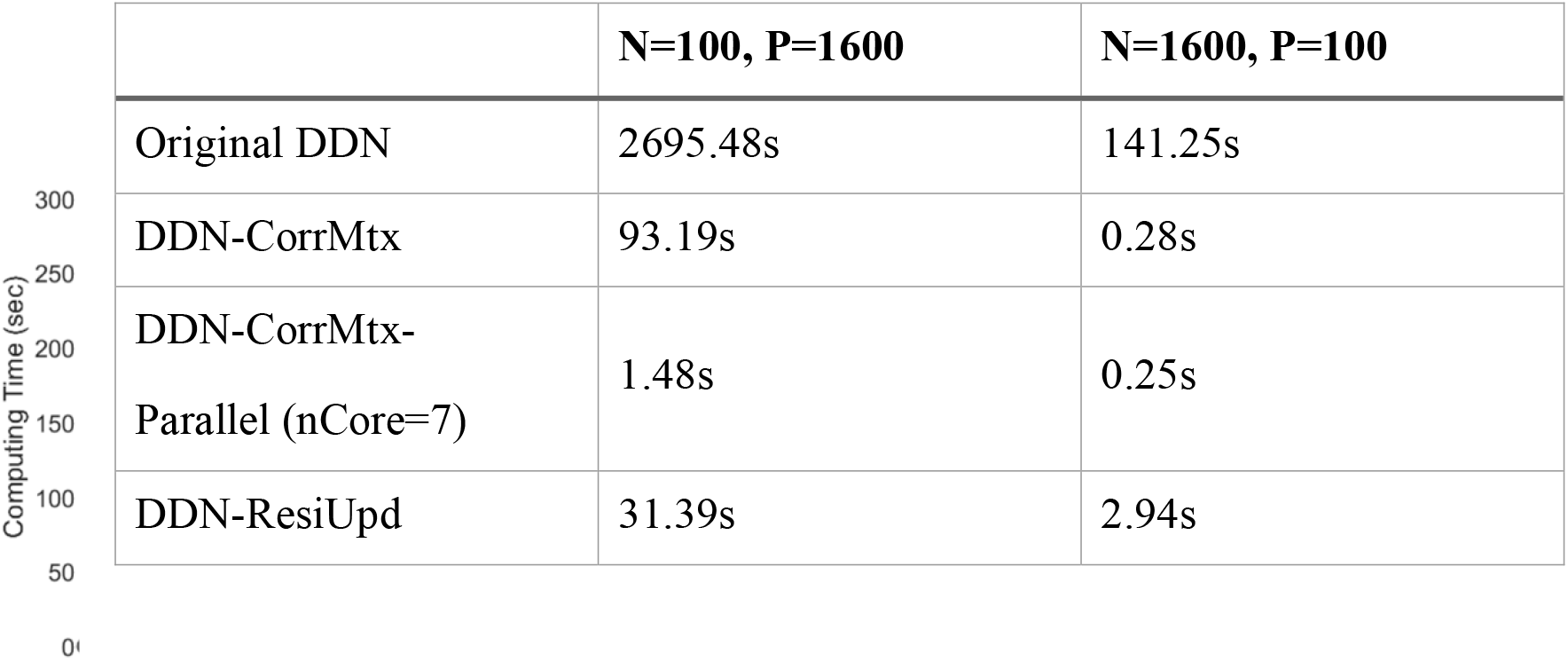
DDN2.0+ computation time versus feature scale P

**Figure 8.**
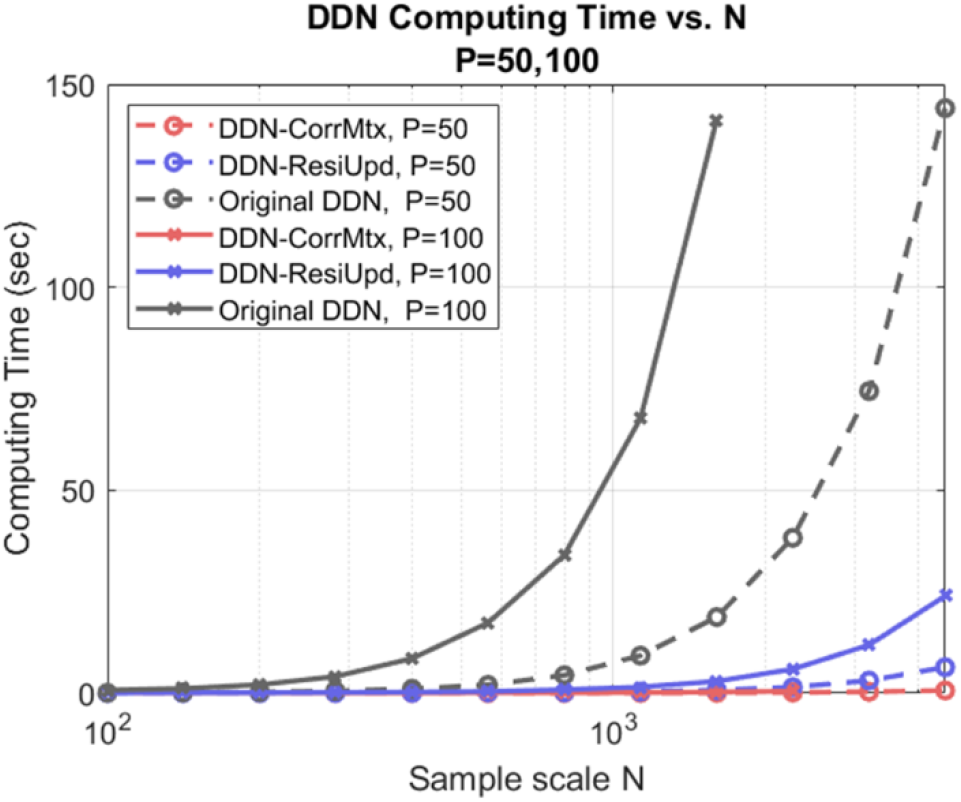
DDN2.0+ computation time versus the sample scale N

**Figure 9.**
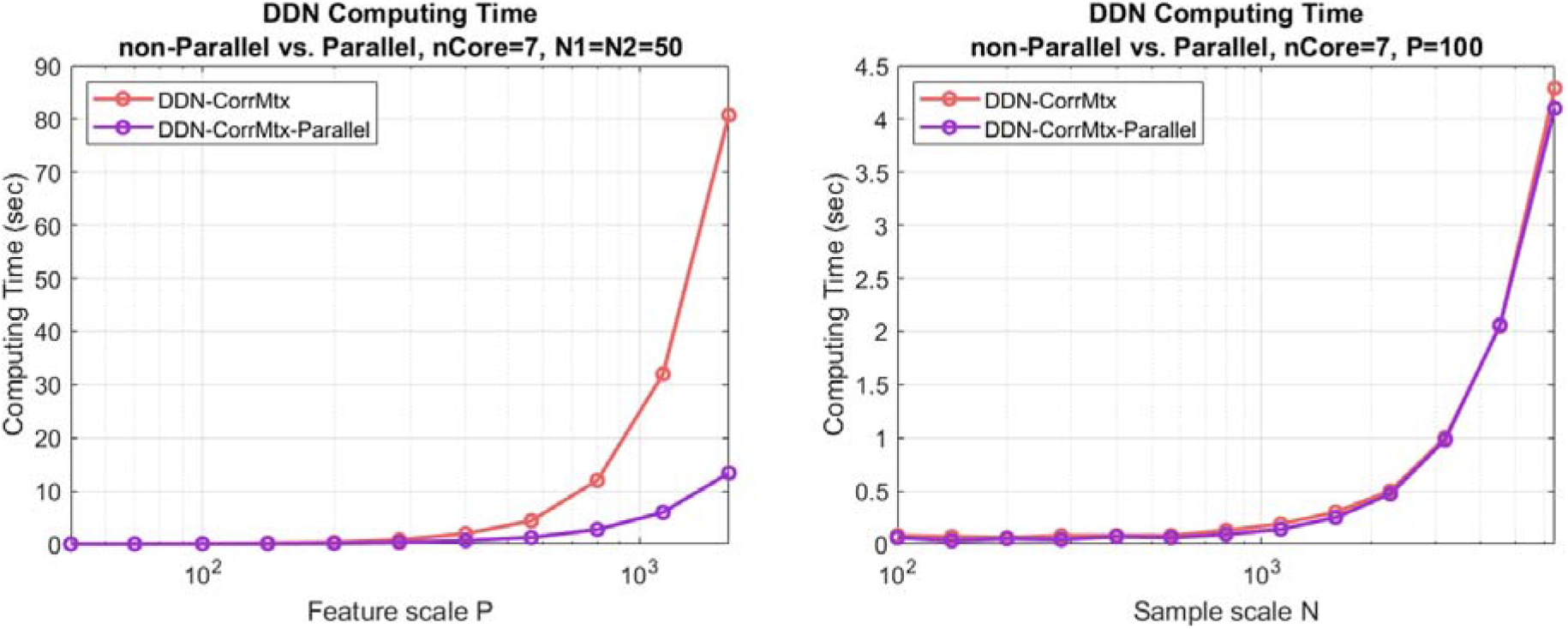
DDN2.0+ computation time comparison between parallel and non-parallel computing

**Figure 10.**
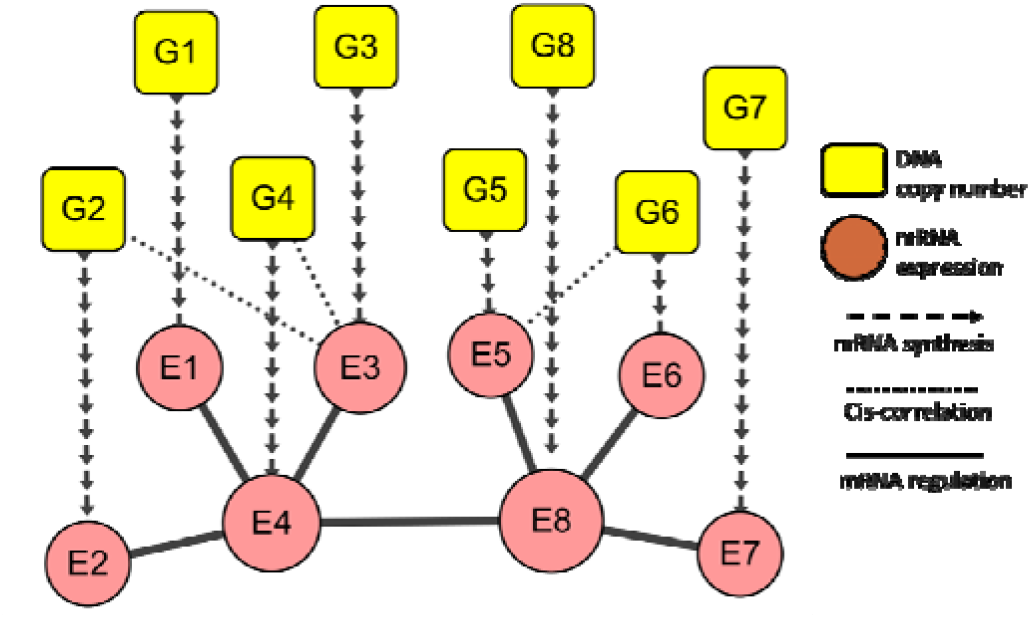
Integrated data model of gene copy number and mRNA expression

**Figure 11.**
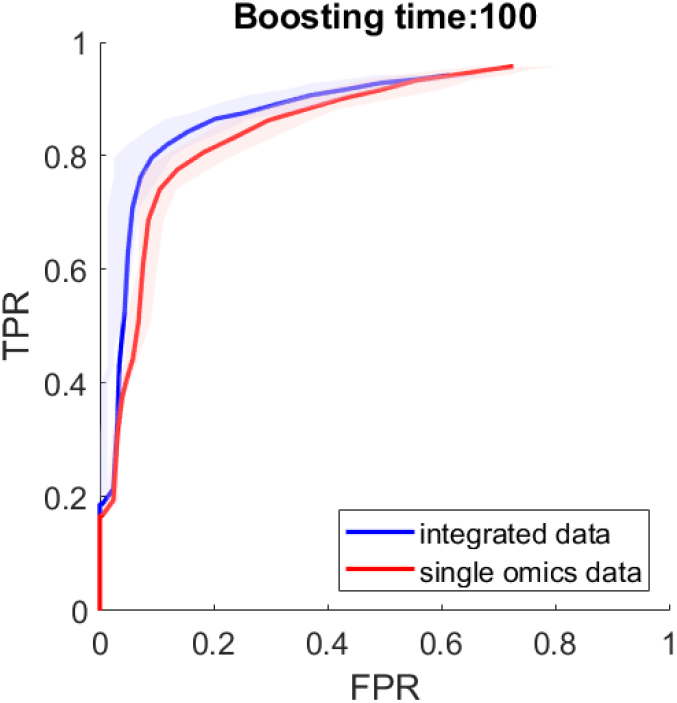
ROC curves of constructing sparse genetic networks from single omics data and from integrated data

**Figure 12.**
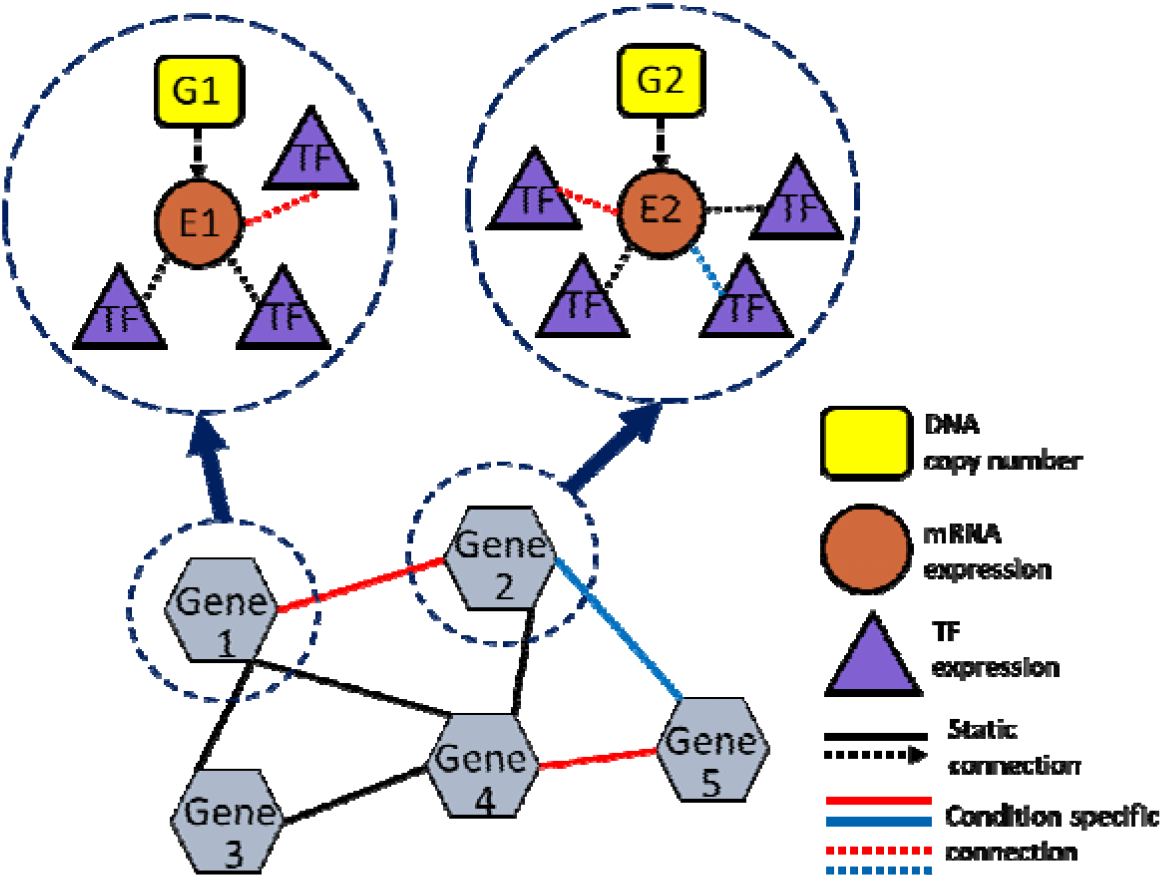
Graphical model of gene entities for multiDDN

**Figure 12.**
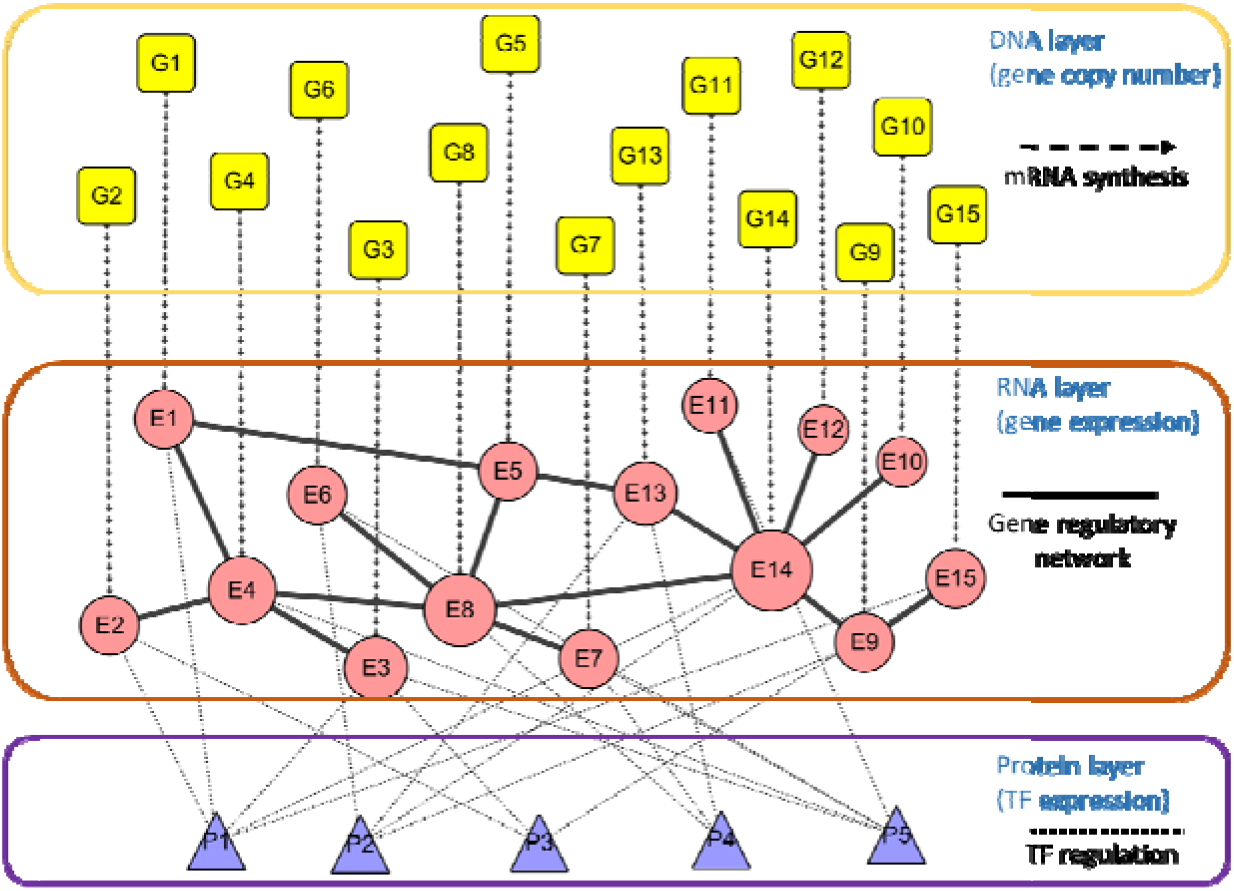
Multi-layer data signaling model for multiDDN

We compare the computation time of the DDN-CorrMtx methods with and without parallel computing (***Figure***). Since parallel computing is done for each node selected from all P nodes, the parallel computing uses much leas computation time when the feature scale P is large. On the other hand, the time saved by parallel computing is limited when P is small.

*Table* shows the effectiveness of the Strong rule. In this simulation study, the ground truth network is designed as a single ring in which each node has exactly two neighbors. Therefore, the basic Strong rules could effectively discard most of the predictors. The results show that computation time could be significantly reduced by applying the Strong rule when the network has high sparsity.

In summary, we proposed three reformulated computing methods for the BCD algorithm used in solving optimization in DDN. We also propose accelerating strategies of discarding predictors by the Strong rule and by parallel computing. Depending on which one of the sample scale N and the feature scale P has the larger value, we may choose from the method of computation with correlation matrix, or residual updating strategy, or with combined effort in coefficient updating strategy. The results show a tremendous reduction in computation time comparing to the original DDN method. The proposed DDN-ResiUpd method with parallel computing is now able to handle hundreds of genes in a reasonable time period (<1 hour), compared with the original DDN method which is only capable of handling dozens of genes.

### 3.3 Multi-omics DDN development

Data integration has a long history in omics data study (Gatza, et al., 2014). There are different categories of data integration methods. While the naïve merging of multi-omics data is simple, it suffers many drawbacks. It completely ignores the inter-omics interaction, and cannot benefit from the knowledge of regulation in the multi-omics data. Some other multi-omics integration methods take the use of inter-omics associations. For example, expression quantitative trait loci (eQTL) analysis uses both genomics and transcriptomics data to detect genomic loci that explain the variation of gene expression levels(Shabalin, 2012). Similarly, methylation quantitative trait loci (mQTL) analysis tries to find loci associated with methylation levels(Volkov, et al., 2016). CPTAC-OV project explored the correlation between chromosome instability summarized from genomics data and the abundance of proteins (Zhang, et al., 2016). These multi-omics integration methods enable researchers to gain unique insights into inter-omics associations.

In integrating multi-omics data in differential networks analysis, we believe that the integration of omics should be more than the sum of its parts. We design a multi-layer integrated signaling model to reduce the total feature scale by incorporating knowledge of inter-omics interaction, accordingly propose a multiDDN that is capable of integrating multi-omics data and detecting both intra- and inter-omics network rewiring.

One of the most import regulatory factors to gene expression level is the gene dosage effect from the transcribed gene itself (Gardiner, 2004). A gene’s dosage which is quantified as the gene copy number determines the maximum number of copies of the gene that can be transcribed into mRNA simultaneously. Yang, et al. (2007) reported that the gene dosage effect could reach as high as r=0.98 in some genes such as HER2 and GRB7. By sorting the significant correlations between gene’s copy number variation and mRNA expression with their genomic locations, we also noticed that gene locations are highly likely to positively correlated with copy numbers from genes in nearby genomic locations. This phenomenon referred to as cis-correlation is likely caused by long-range structural variations in the DNA chain, and is confirmed by many reports (Bryois, et al., 2014; Yang, et al., 2007). It is one of the major confounding factors in gene regulatory network inference, and our integrated model aims to alleviate this confounding factor.

Based on the facts of the gene dosage effect of copy numbers, we propose an integration method to add DNA copy number information as additional predictors to gene expression variation in the gene regulation network construction. ***Figure*** shows an illustrative example of this model shows how the DNA copy number information interacts with the gene regulation network in the mRNA level. Although DNA may carry germline or somatic copy number variations in living cells, it is commonly believed that the numbers of gene copies are not regulation targets and are not altered by other biomolecules. Therefore, in the network construction we may treat DNA copy number signals solely as input predictor variables. We also limit its response variables as its gene expression variable due to the strong gene dosage effect, so that only one additional predictor is added to each node in gene expression layer, and the total feature scale in each node’s DDN optimization problem will be p+1 instead of 2p.

A pilot simulation study is designed to show the integrated model’s effectiveness in inferring a gene regulatory network. We predesigned a gene regulatory network of 15 genes in the RNA layer, and associate the genes’ expression to their own copy numbers in the DNA layer and some additional weaker links that represent the cis-correlation effect. The copy number and gene expression values are sampled from multivariate Gaussian distributions. We compare two methods to reconstruct the sparse networks: the first one is the baseline LASSO regression approach for the single-omics data (gene expression data) alone; and the second one is the integrated method of the LASSO regression with an additional predictor of copy number of the same gene. We use the receiver operating characteristic curve (ROC) which is the curve of the true positive rate (TPR) against the false positive rate (FPR) at various threshold settings. We also use the bootstrapping method with 100 times of boosting to evaluate the confidence interval (CI) of the ROC. The simulation result shows the integrated method has a larger area under the curve (AUC) than the method on single-omics data alone. And for a given false positive rate (FPR), the integrated method has a significantly higher true positive rate than the single-omic method.

The role of TFs in the integrated data signaling model is the regulators of the genes in the RNA layer. Gene regulatory network is inferred mainly from gene expression data and from inter-omics dependency between gene expression and TF expression. We are not detecting regulations between TFs in this model, but the method of multiDDN could be easily extended to include such dependencies between nodes in the protein layer.

There are various ways of selecting candidate nodes from the whole set of molecules in omics data. For multiDDN, we may choose genes from pathways of interest or curated gene list; or if pathway information is not available, we significantly differentially expressed genes from transcriptomics data. We select the TF nodes for the integration model in the following way: The TF-gene binding information is retrieved from the TRRUST database(Han, et al., 2017), restricted to the Homo Sapiens and with experimental evidence. The genes in the RNA layer are used as input to find their regulating TFs. The retrieved list of TFs is further filtered to keep only those regulate at least three genes in the RNA layer and P-value less than 1E-3.

The multi-layer model could be summarized to dependency networks of gene entities. Gene entity in a multiDDN network is defined as one gene’s expression combined with its own copy number regulator and with its regulating TFs. The differential dependency (i.e., the network rewiring) in the multiDDN network is in two tiers: the first is the intra-omics network rewiring between the gene entities; and the second is the inter-omics network rewiring within a gene entity. ***Figure*** illustrates the conceptional multiDDN network between gene entities.

Now consider the problem of learning graphical structure changes in the data model between two conditions. The problem is equivalent to estimate the conditional dependence or independence between a subset of random variables as gene entities under two conditions, with additional variables to each gene entity as entity-specific predictors. We have a set of *p*_*G*_ = *p*_*E*_ = *p* genes of interest which are bind with a total of *P*_*p*_ TF proteins. We observed samples from *n*_1_ objects under condition 1, and *n*_2_ objects under condition 2. For each object, we collected three variables of copy number, gene expression and TF expression.

For convenience, we firstly define a few terms. Define the vectors of variables observed from the *i*-th sample under condition 1 as:

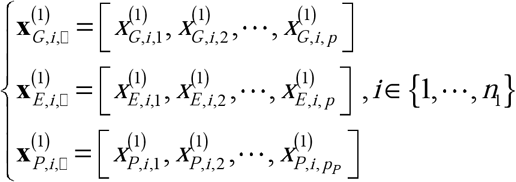

in which the letter G, E, P represents data from genomics data, gene expression data and protein data, respectively. The vectors of variables under condition 2 are defined in a similar manner.

In the dimension of features, denote all observation on *j*-th gene or *j’*-th TF protein under condition 1 as:

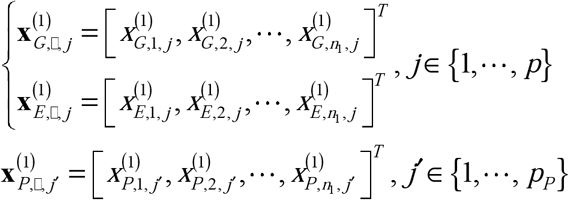

Similarly denote the observation vectors under condition 2. The vectors of variables are merged to data matrices of omics, either from sample dimension or feature dimension.

Define:

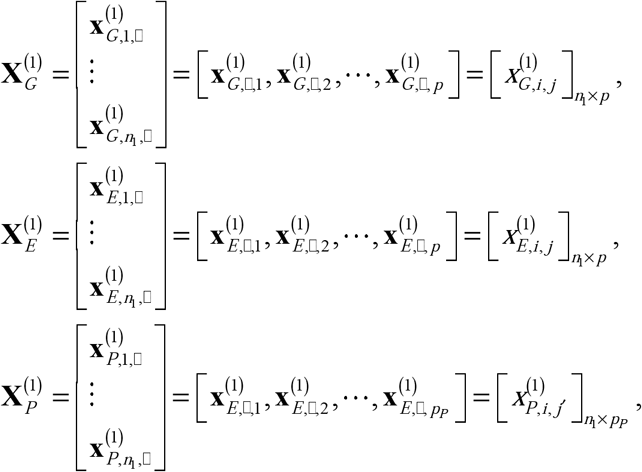

as the three data matrices of three omics under condition 1, and similarly for condition 2. 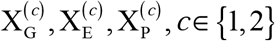 are data matrices of gene copy number, mRNA expression and protein expression, respectively. Denote 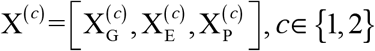 as the entire observation data matrix under conditions 1 or 2. For the *j*-th gene entity, omics under condition 1 or 2, define the coefficient β vector which is combined from three sub β vectors from each omics as:

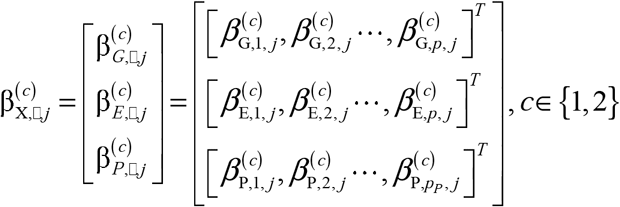

The β vector under condition 1 or 2 for all gene entities are merged into the β matrix which is the representation of the multiDDN network structure:

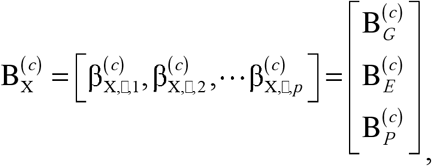

Along the sample feature dimension, the β vector for the *j*-th gene entity under both conditions form the network dependency structure for the *j*-th gene entity, define as:

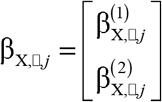

Finally, define two LASSO objective function for each condition and the multiDDN’s objective function as:

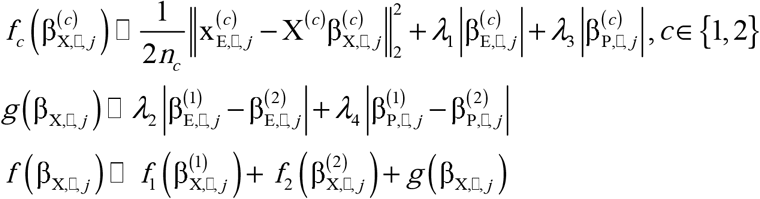

Mathematically, we formulate a multiDDN problem of learning structural changes of the multi-omics data model between two conditions as a convex optimization problem. We solve the following optimization problem for the *j*-th gene entity as follows (*j* = 1, 2, …, *p*):

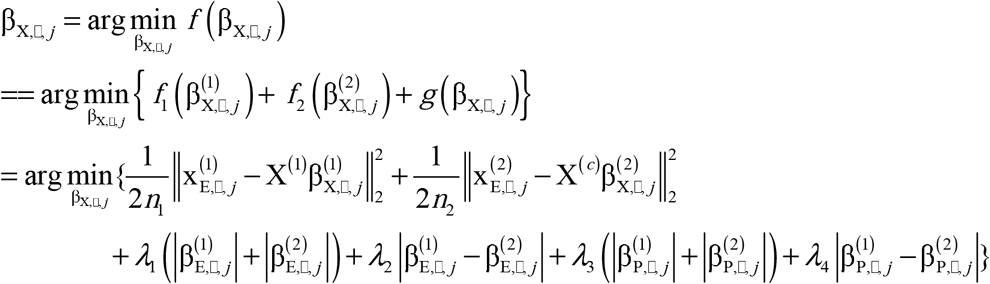

In the multiDDN optimization problem, we learn the graphical structures of the integrated data model under two conditions jointly. The LASSO objective function 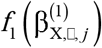 and 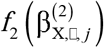 for each condition lead to the identification of a sparse graph structure. The penalty term *g*(β_X,◻,*j*_), encourages sparse changes in the network structure of both intra-omics and inter-omics interactions between two conditions, and thereby suppresses parametric inconsistencies due to noise or limited samples.

After solving the multiDDN optimization problem for each gene entity, the matrix of 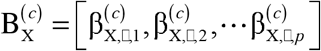 are the parametric representation of the multiDDN network under each condition. For 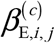 and 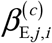, we may replace them with the one with the larger absolute value to get a symmetric parametric structure for the intra-omics part 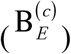 which could be converted to adjacency matrix for nodes in the RNA layer. The two parametric representations 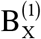 and 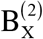 are then compared to exact the network rewiring events from the differential matrix 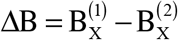. We may further separate the differential matrix to Δβ_*G*_ Δβ_*E*_ and Δβ_*P*_, to categorize the network rewiring events into intra-omics network rewiring and inter-omics network rewiring.

We design a simulation study to show that our proposed multiDDN method will have higher prediction precision in constructing differential networks from the multi-omic data, comparing with the DDN method on single omics data. The ground truth of the multi-layer regulation network is designed as a scale-free network to mimic the gene regulatory network form real biological data, as shown in ***Figure 1***. We then generate the adjacency network according to the ground truth. For the regulation strengths, we measure the intra-omics Pearson’s correlation coefficient distribution from over 200 samples in TCGA ovarian cancer dataset (Cancer Genome Atlas Research, 2011; Zhang, et al., 2016), and take the mean and variance values as the guiding parameters for the simulated covariance matrix, giving the fact that the correlation matrix is identical to the covariance matrix for standardized data. Similarly, we measure inter-omics Pearson’s correlation coefficient distribution for interactions between DNA and mRNA, and interactions between TF with mRNA. Depending on the type, the non-zero elements in simulated covariance Σ are sampled from one of the three distributions, and the whole data matrix is then sampled from multivariate Gaussian distribution of *G* (0, Σ). These steps are repeated once again with a slightly different network skeleton and covariance matrix, to generate the data matrix of condition 2. The network structure differences between conditions 1 and 2 are recorded as the ground truth of network rewiring, as shown in ***Figure 1*** as colored edges.

**Figure 1.**
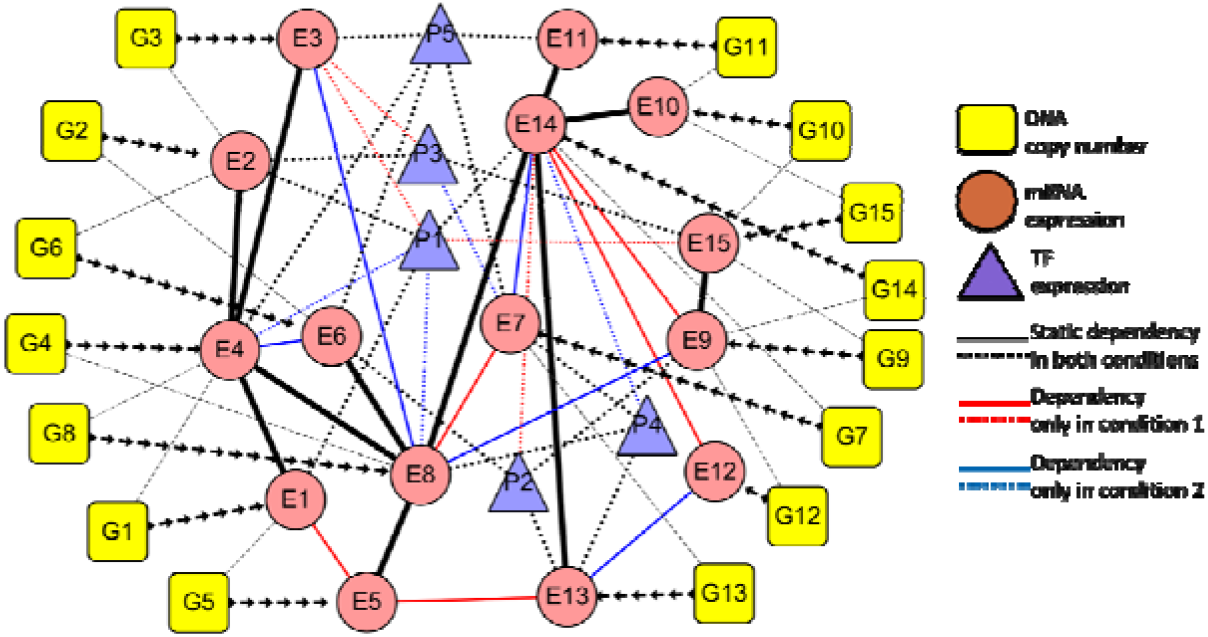
Synthesized multi-layer differential network used in multiDDN simulation

We test the multiDDN method on four groups of simulated data: 1. Single omics data that contains only gene expression; 2. Two-omics data combined from copy number data and gene expression data; 3. Two-omics data combined from gene expression data and TF expression data; 4. Three-omics data combined from all three types of omics data. To compare the results on a fairground, only the common layer of four groups’ networks which is the RNA layer is compared. The multiDDN performance in these four groups is shown in Figure 14 *The ROC curves for multiDDN on multi-omics data with different integration levels. The black curve is for the multiDDN method with all three types of omics data as the input. The blue and red curves are for the multiDDN method with only two of the three types of omics data. The green curve is for the DDN method with single-omics data of mRNA expression.*

As expected, multiDDN on the integrated three omics data has the largest area under the ROC curve, and DDN on single omics data has the smallest. This confirms the benefit of integrating additional omics data into differential network analysis.

**Figure 14.**
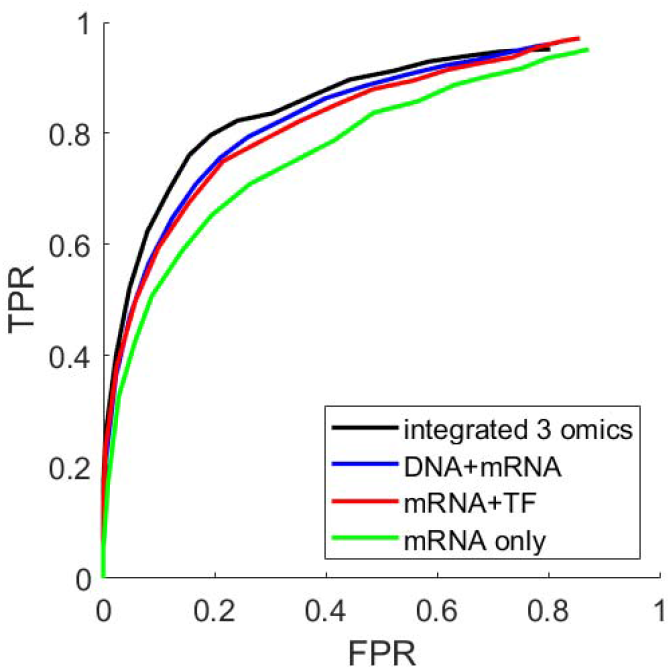
The ROC curves for multiDDN on multi-omics data with different integration levels. The black curve is for the multiDDN method with all three types of omics data as the input. The blue and red curves are for the multiDDN method with only two of the three types of omics data. The green curve is for the DDN method with single-omics data of mRNA expression.

